# Completing bacterial genome assemblies with multiplex MinION sequencing

**DOI:** 10.1101/160614

**Authors:** Ryan R. Wick, Louise M. Judd, Claire L. Gorrie, Kathryn E. Holt

## Abstract

Illumina sequencing platforms have enabled widespread bacterial whole genome sequencing. While Illumina data is appropriate for many analyses, its short read length limits its ability to resolve genomic structure. This has major implications for tracking the spread of mobile genetic elements, including those which carry antimicrobial resistance determinants. Fully resolving a bacterial genome requires long-read sequencing such as those generated by Oxford Nanopore Technologies (ONT) platforms. Here we describe our use of the ONT MinION to sequence 12 isolates of *Klebsiella pneumoniae* on a single flow cell. We assembled each genome using a combination of ONT reads and previously available Illumina reads, and little to no manual intervention was needed to achieve fully resolved assemblies using the Unicycler hybrid assembler. Assembling only ONT reads with Canu was less effective, resulting in fewer resolved genomes and higher error rates even following error correction with Nanopolish. We demonstrate that multiplexed ONT sequencing is a valuable tool for high-throughput bacterial genome finishing. Specifically, we advocate the use of Illumina sequencing as a first analysis step, followed by ONT reads as needed to resolve genomic structure.

**Data summary:**

1. Sequence read files for all 12 isolates have been deposited in SRA, accessible through these NCBI BioSample accession numbers: SAMEA3357010, SAMEA3357043, SAMN07211279, SAMN07211280, SAMEA3357223, SAMEA3357193, SAMEA3357346, SAMEA3357374, SAMEA3357320, SAMN07211281, SAMN07211282, SAMEA3357405.
2. A full list of SRA run accession numbers (both Illumina reads and ONT reads) for these samples are available in **Table S1**.
3. Assemblies and sequencing reads corresponding to each stage of processing and analysis are provided in the following figshare project: https://figshare.com/projects/Completing_bacterial_genome_assemblies_with_multiplex_MinION_sequencing/23068
4. Source code is provided in the following public GitHub repositories: https://github.com/rrwick/Bacterial-genome-assemblies-with-multiplex-MinION-sequencing https://github.com/rrwick/Porechop https://github.com/rrwick/Fast5-to-Fastq

**Impact Statement:** Like many research and public health laboratories, we frequently perform large-scale bacterial comparative genomics studies using Illumina sequencing, which assays gene content and provides the high-confidence variant calls needed for phylogenomics and transmission studies. However, problems often arise with resolving genome assemblies, particularly around regions that matter most to our research, such as mobile genetic elements encoding antibiotic resistance or virulence genes. These complexities can often be resolved by long sequence reads generated with PacBio or Oxford Nanopore Technologies (ONT) platforms. While effective, this has proven difficult to scale, due to the relatively high costs of generating long reads and the manual intervention required for assembly. Here we demonstrate the use of barcoded ONT libraries sequenced in multiplex on a single ONT MinION flow cell, coupled with hybrid assembly using Unicycler, to resolve 12 large bacterial genomes. Minor manual intervention was required to fully resolve small plasmids in five isolates, which we found to be underrepresented in ONT data. Cost per sample for the ONT sequencing was equivalent to Illumina sequencing, and there is potential for significant savings by multiplexing more samples on the ONT run. This approach paves the way for high-throughput and cost-effective generation of completely resolved bacterial genomes to become widely accessible.

## Introduction

The low cost and high accuracy of Illumina sequencing reads makes them well suited to high-throughput bacterial genomics. Read-mapping pipelines can use Illumina reads to reliably identify single nucleotide polymorphisms, necessary for inferring phylogenies and understanding population structure [1]. Illumina reads can quickly reveal a sample’s sequence type and whether genes of clinical interest are present in the genome [2]. This has caused Illumina platforms to become the dominant technology for whole genome sequencing of bacterial isolates, including for routine public health applications, with hundreds of thousands of Illumina read sets and genome assemblies now publicly available [3]. The major shortcoming of Illumina reads is their length: maximally 300 bp but more commonly ≤150 bp. Short reads cannot resolve all genomic repeats, and fragmented assemblies comprising hundreds of discrete contigs are often the best possible outcome [4]. Of particular concern to many, antimicrobial resistance regions are often flanked by repetitive insertion sequences, and it can be impossible to tell from an incomplete short-read assembly whether genes of interest reside in the chromosome or on a plasmid [5 – 7]. This is a hindrance to researchers and public health laboratories, as the location of resistance genes can have significant epidemiological implications [8,9].

Alternative sequencing platforms produced by Oxford Nanopore Technologies (ONT) and Pacific Biosciences (PacBio) can generate “long reads” that are several kilobases in length. These reads can exceed the length of repeats in a typical bacterial genome, making complete assembly (with one contig per replicon) possible [10,11]. With long-read assemblies, the full structure of a bacterial genome comes into focus, revealing the position of all genes. Despite this key advantage, the accuracy of the resulting sequence varies, even after using error-correcting tools such as Nanopolish [12].

Due to the popularity of Illumina sequencing and the relative ease of generating long reads using the cheap and portable ONT MinION sequencer, there is increasing interest in “hybrid” assemblies that combine both types of data to produce complete genome assemblies that are highly accurate in both structure and sequence accuracy. The hybrid approach suits the common work flow used by a wide range of research and public health laboratories, whereby new isolates are sequenced *en masse* with Illumina to facilitate species identification, gene profiling and strain relatedness or transmission patterns. Long reads can then be added where needed to resolve plasmids and complex antibiotic resistance or phage regions via hybrid assembly [3]. To facilitate this approach, we recently developed a software tool, Unicycler, which can automatically generate high-quality hybrid assemblies of complete bacterial genomes using a combination of Illumina and long-read data [13].

Here we describe a protocol for the generation of complete, highly accurate bacterial genome sequences at relatively high-throughput and low cost, using the ONT barcoding kit to generate long reads for 12 isolates simultaneously on a single MinION flow cell, followed by hybrid assembly with Unicycler. To demonstrate the utility of the approach, we use it to finish the genomes of 12 clinical isolates of *Klebsiella pneumoniae*. These isolates were previously sequenced with Illumina platforms, but the resulting draft assemblies were inadequate for resolving the position of antimicrobial resistance genes [14]. The total cost of ONT sequencing (reagents and flow cell) for 12 bacterial isolates was approximately 950 USD, or 80 USD per sample, comparable to per-sample costs for Illumina.

## Methods

The scripts used to perform all bioinformatics analyses are available at: https://github.com/rrwick/Bacterial-genome-assemblies-with-multiplex-MinION-sequencing

### Bacterial isolates and Illumina sequencing

*K. pneumoniae* were isolated from patients in a Melbourne hospital as part of a prospective longitudinal study on *K. pneumoniae* carriage and infection [14]. Illumina sequence data for all 12 isolates was generated previously from DNA extracted using a phenol:chloroform protocol [14]. Illumina data was generated using either Nextera libraries followed by Illumina HiSeq 2500 sequencing (at the Australian Genome Research Facility, n=4 isolates) or TruSeq libraries followed by Illumina HiSeq 2000 sequencing (at the Wellcome Trust Sanger Institute, n=8 isolates) [14]. See **Table S1** for properties and accession numbers for the Illumina read sets.

### DNA isolation for ONT libraries

Clinical isolates were grown overnight at 37°C on LB Agar plates, then single colonies picked for overnight culture at 37°C in LB. Bacterial cell pellets from 3.0 ml of LB culture were generated by centrifugation at 15,000 *g* for 5 minutes. DNA was extracted from these pellets using Agencourt GenFind V2 (Beckman Coulter) with minor modifications as follows. Cell pellets were resuspended in 400 μl lysis buffer containing 9 μl Proteinase K (96 mg / ml Beckman Coulter) and 1 μl RNase A (100 mg/ml Sigma Aldrich R6513) by gentle tip mixing. Samples were lysed at 37°C for 30 minutes. We extracted gDNA from the lysed samples by completing the remaining steps of the GenFind V2 for 200 μl of blood / serum from the binding step on. This extraction protocol generates high molecular weight gDNA (>60 kbp) free of small DNA contamination that is suited to ONT MinION sequencing without further purification or size selection.

### ONT library preparation

Libraries were prepared without shearing to maximise sequencing read length. Data yield from ONT MinION sequencing is correlated with number of DNA molecules in the library tagged with adapter allowing entry into nanopores. Working with high molecular weight (unsheared) DNA reduces the number of DNA ends available for adapter ligation so we made efforts to maximise DNA quantity throughout the library preparation.

The library was prepared using the ONT 1D ligation sequencing kit (SQK-LSK108) with the native barcoding expansion kit (EXP-NBD103). The ONT protocol for native barcoding genomic DNA sequencing was followed with the following modifications to maximise DNA recovery. At least 1 μg DNA from each isolate was treated with the end-repair / dA tailing module but the DNA was eluted in 24 μl following AMPure XP bead clean up. Following the barcode ligation reaction, the DNA was cleaned again with AMPure XP beads and elution in 10 μl. For library pooling, we calculated the amount of DNA to add based on 5 μl of DNA from the sample with the lowest concentration. All other samples were added accordingly to produce an equimass pool, and we used 50 μl of this pooled DNA sample for adapter ligation.

### MinION Sequencing

The final library containing 2415 ng DNA was loaded onto an R9.4 flow cell. The run was performed on a MinION MK1b device using the NC_48Hr_Sequencing_Run_FLO-MIN106_SQK-LSK108 protocol with 1432 available pores (508, 459, 337 and 128 pores per group). The protocol terminated prematurely after 7.25 hours due to a MinKNOW software crash. At this point we started the run again and it proceeded for the full 48 hours. All data described is the accumulation of the two runs.

### Basecalling and read preparation

After the sequencing run finished, we transferred the fast5 read files to a separate Linux server and performed basecalling with ONT’s Albacore command line tool (v1.1.2). Albacore was run with barcode demultiplexing and fastq output. We then trimmed adapter sequences from the reads using Porechop (v0.2.1, https://github.com/rrwick/Porechop). To reduce the risk of cross-barcode contamination, we ran Porechop with barcode demultiplexing and only kept reads for which Albacore and Porechop agreed on the barcode bin. The resulting 12 demultiplexed, barcode-trimmed read sets were deposited in under accessions SRR5665590 – SRR5665601 (**Table S1**) and plotted in **Fig 1 and S1**.

**Figure 1:**
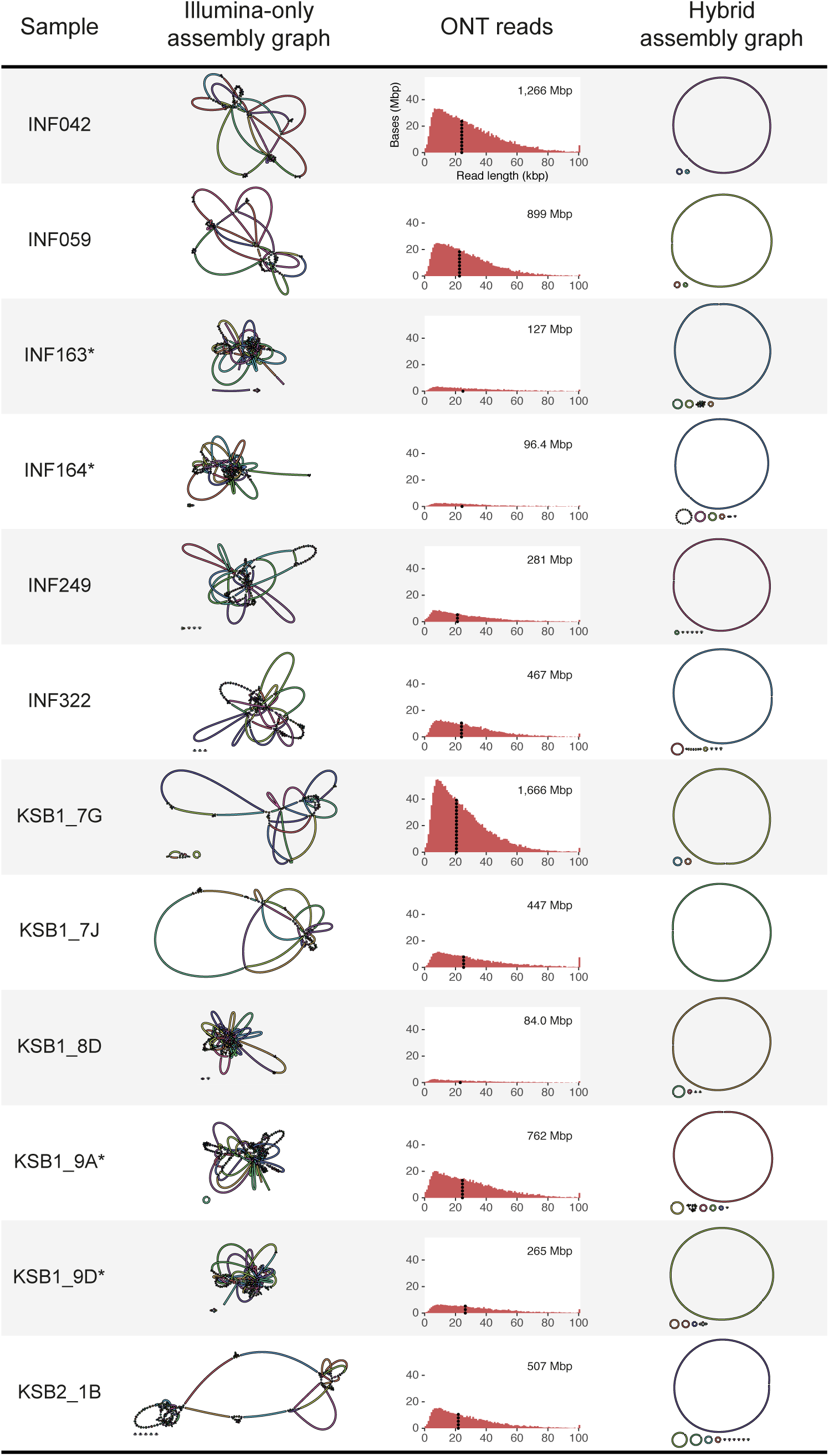
Assembly graphs and length distribution of ONT reads are shown for each *K. pneumoniae* isolate. Both Illumina-only and hybrid assembly graphs were produced with Unicycler (v0.4.0). ONT total yield for each sample is shown in the top right of each histogram. Read N50 length is indicated on each histogram with a dotted line. * Illumina library was prepared using Nextera (all others used TruSeq).

We then subsampled each read set down to 500 Mbp high quality reads using an in-house script (https://github.com/rrwick/Fast5-to-Fastq/blob/master/fastq_to_fastq.py). First, reads <2 kbp in length were excluded. If the total read length still exceeded 500 Mbp per sample, reads with low-quality regions (as measured by mean Phred quality scores over a 100 bp window) were excluded until only 500 Mbp of sequence remained. This ensured that for read sets with abundant sequence, only higher quality reads were used for assembly. The subsampled read sets are available on figshare (https://figshare.com/articles/Subsampled_ONT_reads/5171491).

### Assembly

We assembled each isolate’s hybrid read set (both Illumina and ONT reads) using Unicycler (v0.4.0) [13]. Briefly, Unicycler performs a SPAdes assembly of the Illumina reads and then scaffolds the assembly graph using long reads. Unicycler polishes its final assembly with Illumina reads and Pilon to reduce the rate of small base-level errors [15]. If Unicycler was unable to produce a complete assembly (i.e. one circular contig per replicon), we manually finalised the assembly as follows. We used minimap to select ONT reads which corresponded to incomplete regions of the assembly, aligned these reads to the assembly graph using Bandage and BLAST, and chose the resolved assembly most compatible with the alignments [16,17].

To allow comparison of our hybrid approach with ONT-only assembly (**Fig S2**), we assembled the subsampled ONT read sets using Canu (v1.5) [11]. Each resulting contig set was then processed with one round of Nanopolish (v0.7.0) to calculate consensus base calls using signal-level analysis (based on the fast5 files) [12]. For Nanopolish we used the full set of ONT reads before subsampling.

### Error rate estimation

To assess the consensus base-call accuracy of our assemblies, we compared them to highly reliable reference contig sequences produced via the following method. Each sample’s Illumina reads were assembled using both ABySS (v2.0.1) and Velvet (v1.2.10) – independent alternatives to Unicycler and Canu [18,19]. To exclude ambiguous parts of the assemblies, we used MUMmer to extract sequences of ≥10 kbp in length for which the ABySS and Velvet assemblies were in exact agreement [20]. The resulting contigs (see data summary for availability) were assumed to be 100% accurate and used as references. We then aligned each assembly’s contigs to the corresponding reference contigs using BLASTN [21]. Only the best alignment was kept for each reference sequence, and the number of single base changes and small indels were counted. The final estimates for the frequency of these base-level errors in each assembly were expressed as a weighted mean of error rates in the resulting alignments (**Table S1**, **Fig S2 and S3**).

### Read depth estimation

To estimate the read depth per replicon, we aligned Illumina and ONT reads to the completed genomes using Bowtie 2 (v2.3.0) and BWA MEM (v 0.7.15-r1140), respectively [22,23]. Samtools depth was used to gather the depth for each base of the assembly, and we took the mean value for each replicon [24]. We then normalised each replicon’s depth against the chromosomal depth of the same read set to generate the plot in **Fig S4**.

## Results

### Sequencing

The ONT MinION sequencing run generated 837,413 reads, totalling 10.48 Gbp after basecalling. 331,272 reads (3.61 Gbp) could not be confidently assigned to a barcode bin, leaving 506,141 reads (6.87 Gbp) available for assembly. The distribution of reads across the 12 bins was uneven, ranging from 92 Mbp (approximately 17x depth for a *Klebsiella* genome) to 1.7 Gbp (approximately 310x depth) (**Fig 1 and S1**). After adapter trimming and barcode binning, reads had a mean length of 13,568 bp and an N50 length of 22,901 bp. Sample-level read and assembly statistics are given in **Fig 1 and Table S1**.

### Hybrid assembly

The Illumina-only Unicycler assemblies showed the Illumina reads to be of high quality (**Fig 1 and Table S1**). Many assembly graphs contained no dead ends, suggesting complete Illumina-read coverage of the genome. However, the four read sets generated from Nextera libraries tended to contain more dead ends than those generated from TruSeq libraries (median 2 vs 0, see **Fig 1 and Table S1**).

For all 12 samples, the Unicycler hybrid assembly produced a complete circular chromosome sequence of near-perfect sequence accuracy (**Fig 1 and Table S1**). For seven samples, all plasmid replicons were also complete and no further work was required. For the remaining five samples (INF163, INF164, INF322, KSB1_9A and KSB1_9D), the Unicycler assembly had unresolved repeats in small plasmid sequences which we could manually resolve (see **Methods**). This may be explained by underrepresentation of small plasmids in ONT reads: while ONT and Illumina read depths were in close agreement for large plasmids (>67 kbp), small plasmids (<7 kbp) sequenced tens to hundreds of times deeper in Illumina reads than in ONT reads, relative to chromosome sequences (**Fig S4**).

One hybrid assembly (INF164) contained an 18 kbp element that was present in the Illumina reads but not in the ONT reads. We produced a manually-completed assembly that is compatible with the ONT reads (which do not include the 18 kbp element) but could not produce an assembly compatible with the Illumina reads (which included the 18 kbp element).

### Long-read-only assembly

ONT-only assembly with Canu produced complete chromosome sequences for five genomes, including four out of five read sets that exceeded 500 Mbp and a single smaller read set of 127 Mbp (**Fig S2 and Table S1**). Large plasmids were usually complete in these assemblies, but all small plasmids were missing, consistent with the low representation of small plasmids in our ONT data sets as outlined above (**Fig S4**). The ONT-only assemblies had an average consensus base-call error rate of 1.22% before Nanopolish and 0.67% after one round of Nanopolish (calculated in non-repetitive sequences only, see **Methods**) (**Fig S2 and S3**).

## Discussion

### Lessons learned

ONT’s library preparation protocols specify quantity of DNA using mass units. However, the chemistry and subsequent sequencing yields do not rely on the mass of DNA but rather on the number of DNA ends (i.e. molarity) which is proportional to mass and inversely proportional to fragment length. ONT’s protocols involve shearing DNA to 8 kbp and the instructions assume this length, but the mean fragment length for unsheared gDNA can be considerably longer. Following the protocol’s mass instructions will therefore underload the flow cells and compromise sequencing yield. A larger input mass is necessary to achieve the molarity assumed in the protocol. It is also useful to assess the size distribution of DNA using a fragment analyser or equivalent. Even a small amount of fragmented DNA can compromise sequence output by reducing the mean fragment size.

Even when attempting to balance DNA input for each barcode, we found it difficult to achieve similar yield for each barcode bin. There was an order-of-magnitude difference between our lowest and highest barcode bin read depths, and the cause of this variation is not clear. However, the fact that Unicycler completed the assembly for the lowest depth sample (KSB1_8D) demonstrates that an even barcode distribution is not necessarily required when performing hybrid assemblies.

Hybrid assembly of a bacterial genome can be approached in several ways. Unicycler employs a short-read-first method, performing an Illumina read assembly and then scaffolding with long reads. An alternative would be to assembly long reads first (e.g. with Canu) and error correct the assembly with short reads [25]. When both Illumina and long-read sets are of high quality, either approach is viable, but we preferred Unicycler for our analyses as it copes with limited amounts of long reads [13]. Furthermore, hybrid Unicycler assemblies can recover small plasmid sequences which are absent in long-read Canu assemblies [25]. However, a long-read-first assembly may be more appropriate in cases where long reads are abundant and Illumina reads are sparse.

Sample INF164 illustrated a particular challenge in hybrid assembly: biological differences between the two read sets, which can arise when DNA is extracted from different subcultures of the same isolate. To avoid this complication and for optimal hybrid assembly in high-throughput, we recommend extracting and storing sufficient DNA for both Illumina and ONT sequencing during the first analysis.

Chimeric reads, two or more separate pieces of DNA joined together in a single read, are known to occur in ONT sequencing and have the potential to give misleading information about genomic structure [26]. We did observe chimeras in our read sets, though some were removed during read trimming with Porechop. Since Unicycler uses ONT reads to scaffold an Illumina graph, it is not sensitive to the presence of chimeras and our pipeline was not affected. We hypothesise that chimeric reads are more common in preparations that involve ligation (e.g. barcode preparation) and less common in ligation-free preparations.

### Are we ready for ONT-only assemblies?

ONT-only assemblies suffered from high error rates, and while high ONT read depth did improve sequence accuracy, the error rate remained substantial for our highest depth samples (**Fig S3**). This suggests that ONT reads currently contain systematic errors for which increased sequencing depth cannot compensate. Our most accurate ONT-only assembly (sample INF042) had an estimated 0.349% error rate after Nanopolish, equivalent to one error per 287 bp, sufficiently high that most 1 kbp genes will contain an error. Such assemblies are unsuitable for multi-locus sequence typing, resistance allele typing, phylogenomics or transmission studies; hence high-accuracy Illumina data is still required for such applications. The lack of representation of small plasmids in ONT data is also potentially concerning for some applications.

### Future considerations

For assembly of *K. pneumoniae* genomes, we found our ONT read lengths (N50 >20 kbp) and depths (>14x) were sufficient to resolve hybrid assemblies. However, other species may have different requirements determined by the frequency and size of their repetitive elements. For example, *Shigella* genomes contain insertion sequences at very high copy number and we have previously demonstrated that they require approximately twice the ONT sequencing depth relative to *K. pneumoniae* to generate finished hybrid assemblies [13,27]. *Acinetobacter* genomes contain a highly repetitive biofilm-associated gene which varies in length but can exceed 25 kbp (e.g. locus ABA1_02934 in *A. baumannii* strain A1) [28]. Assembly of *Acinetobacter* will therefore be particularly dependent read length to ensure that this repetitive region is spanned by long reads [13].

The underrepresentation of small plasmids in our ONT reads (**Fig S4**) was the primary reason Unicycler failed to automatically produce a completed assembly. We have two hypotheses for this underrepresentation: 1) the DNA extraction method is optimised for large DNA fragments; and 2) our omission of a DNA shearing step during library preparation left small plasmids in a circular state and with no ends available to ONT adapters. It may be worth exploring alternative DNA extraction and shearing methods, especially when numerous small plasmids are expected in the genome.

While our error rate estimates for hybrid assemblies were very low, they only accounted for non-repetitive regions of the genome. Polishing an assembly relies on read alignment, but short reads can suffer from non-specific alignment in repeat regions [29]. Most base-level errors which remain in an Illumina-polished assembly (such as those from Unicycler) are therefore likely to be in genomic repeats such as the RNA operon [30].

## Conclusions

In this study, we used hybrid read sets to produce finished assemblies for 12 genomes for a cost of ∼ $USD150 per strain and with minimal bioinformatics effort. While it was feasible to assemble ONT reads alone, doing so compromised sequence accuracy and recovery of small plasmids. Future improvements to library preparation and base-calling methods may mitigate these issues, but until then both Illumina and long reads are needed to produce ideal assemblies, which we have shown can be achieved in a cost effective and high-throughput manner.

## Author statements

This work was funded by the NHMRC of Australia (fellowship #1061409 to KEH).

The authors declare that there is no conflict of interest regarding the publication of this article.

---

**Figure S1:**
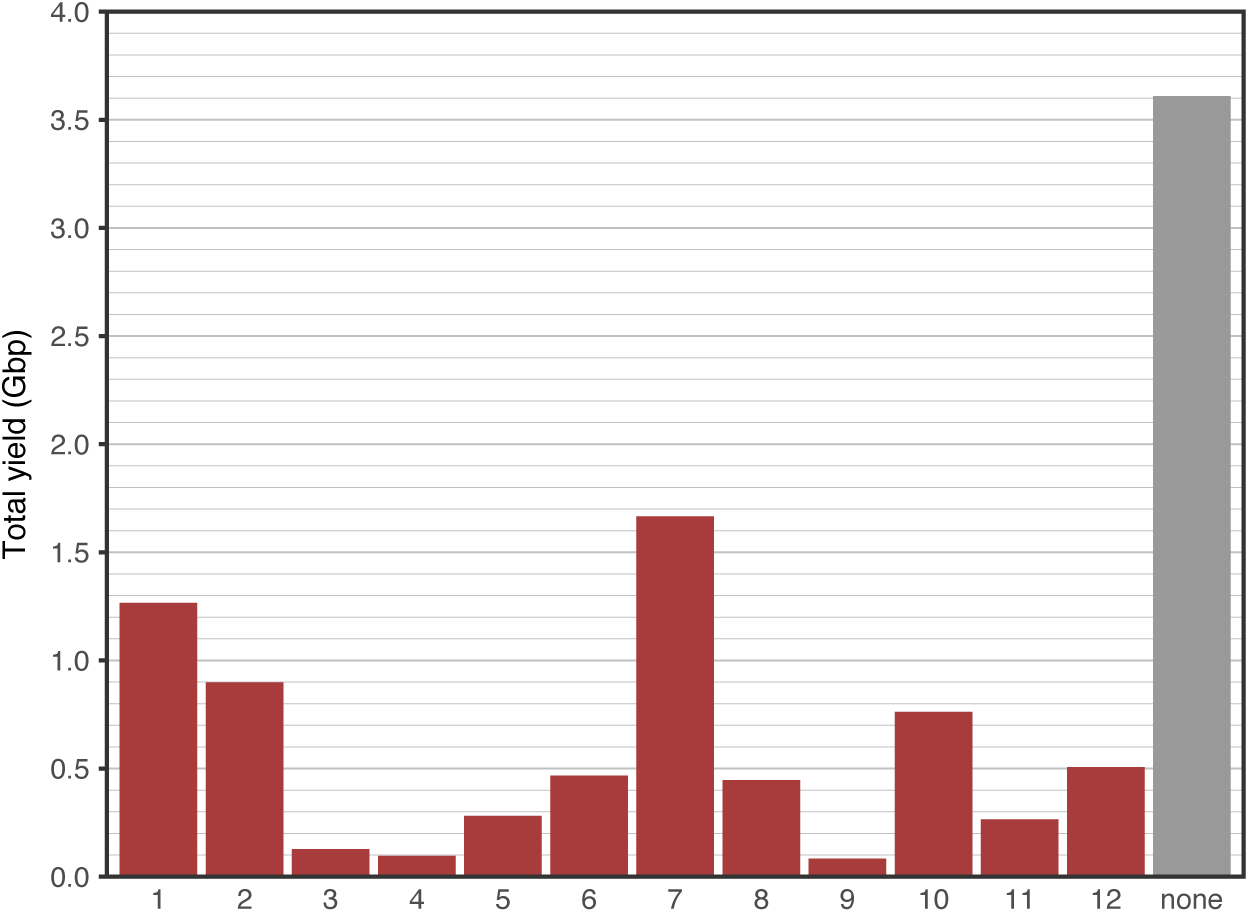
ONT sequencing yield per barcode bin.

**Figure S2:**
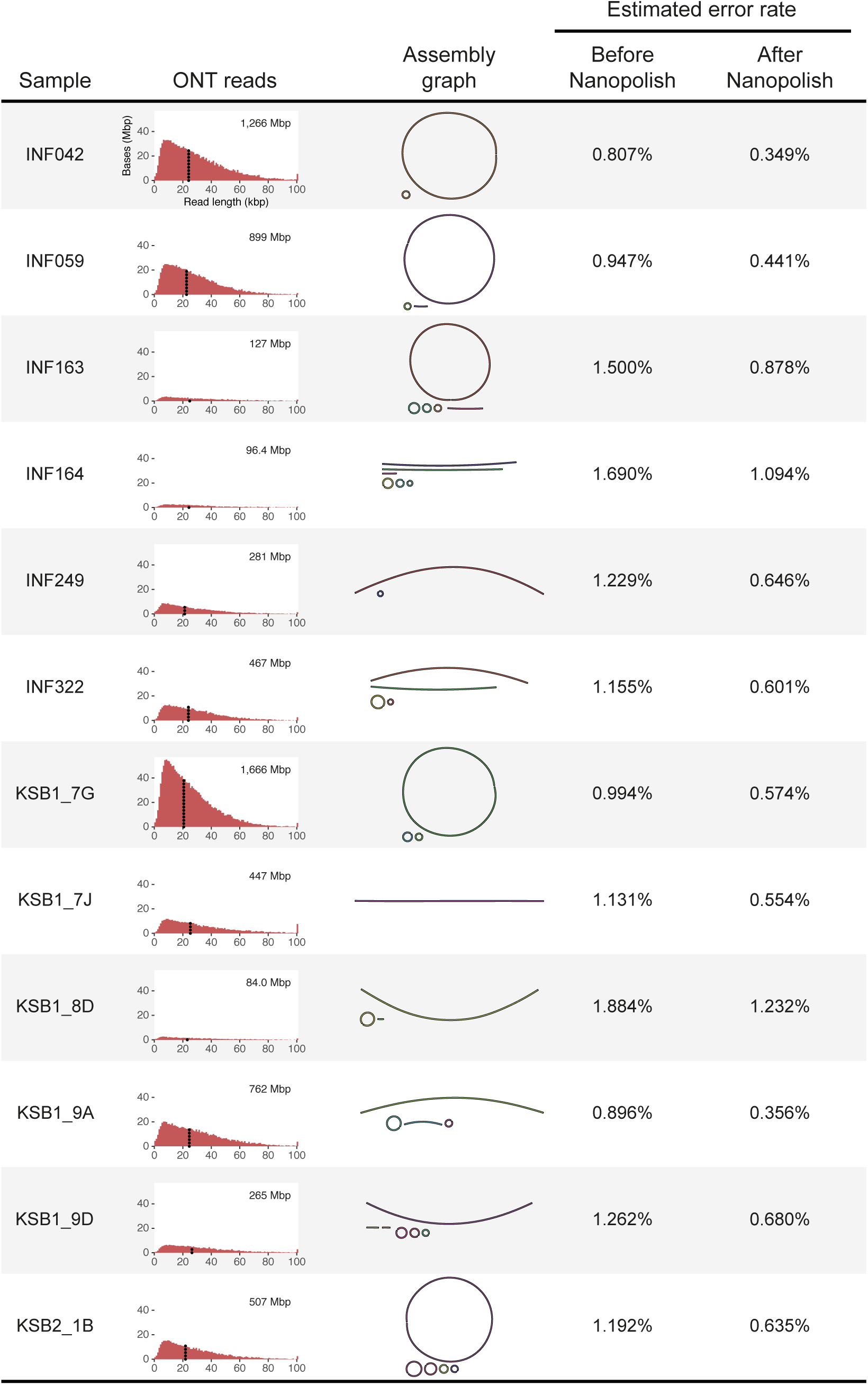
Assembly graphs and error rates for ONT-read-only assemblies produced using Canu (1.5). ONT-read length distributions are shown as histograms and labelled with total ONT yield and N50 read length (reproduced from **Fig 1**). Error rates for each assembly (estimated for non-repetitive regions only, see **Methods**) are shown before and after signal-level consensus base-calling with Nanopolish (v0.7.0).

**Figure S3:**
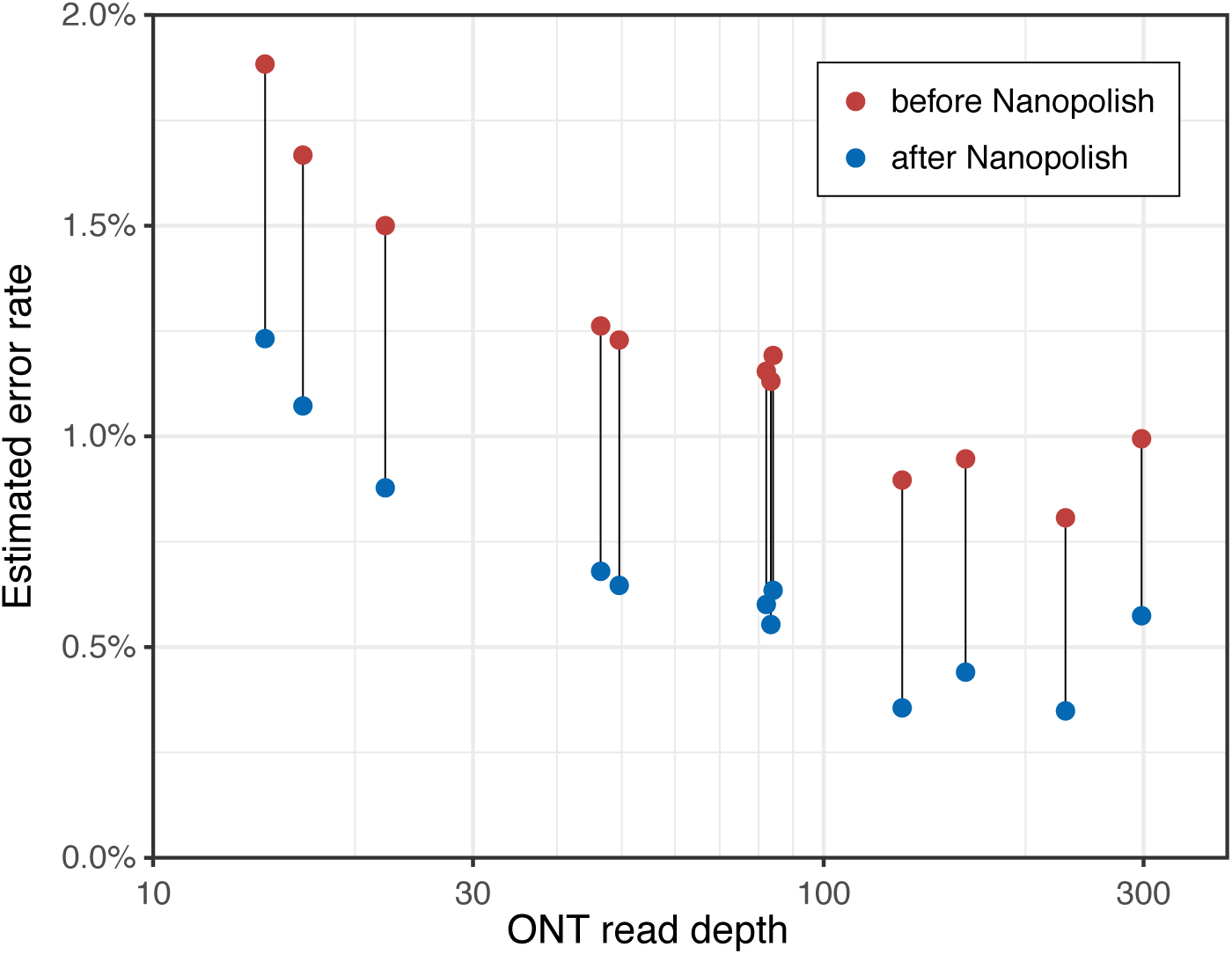
Estimated error-rates in ONT-only Canu assemblies by read depth. Each sample shows the error-rate (in non-repetitive sequences) before and after Nanopolish.

**Figure S4:**
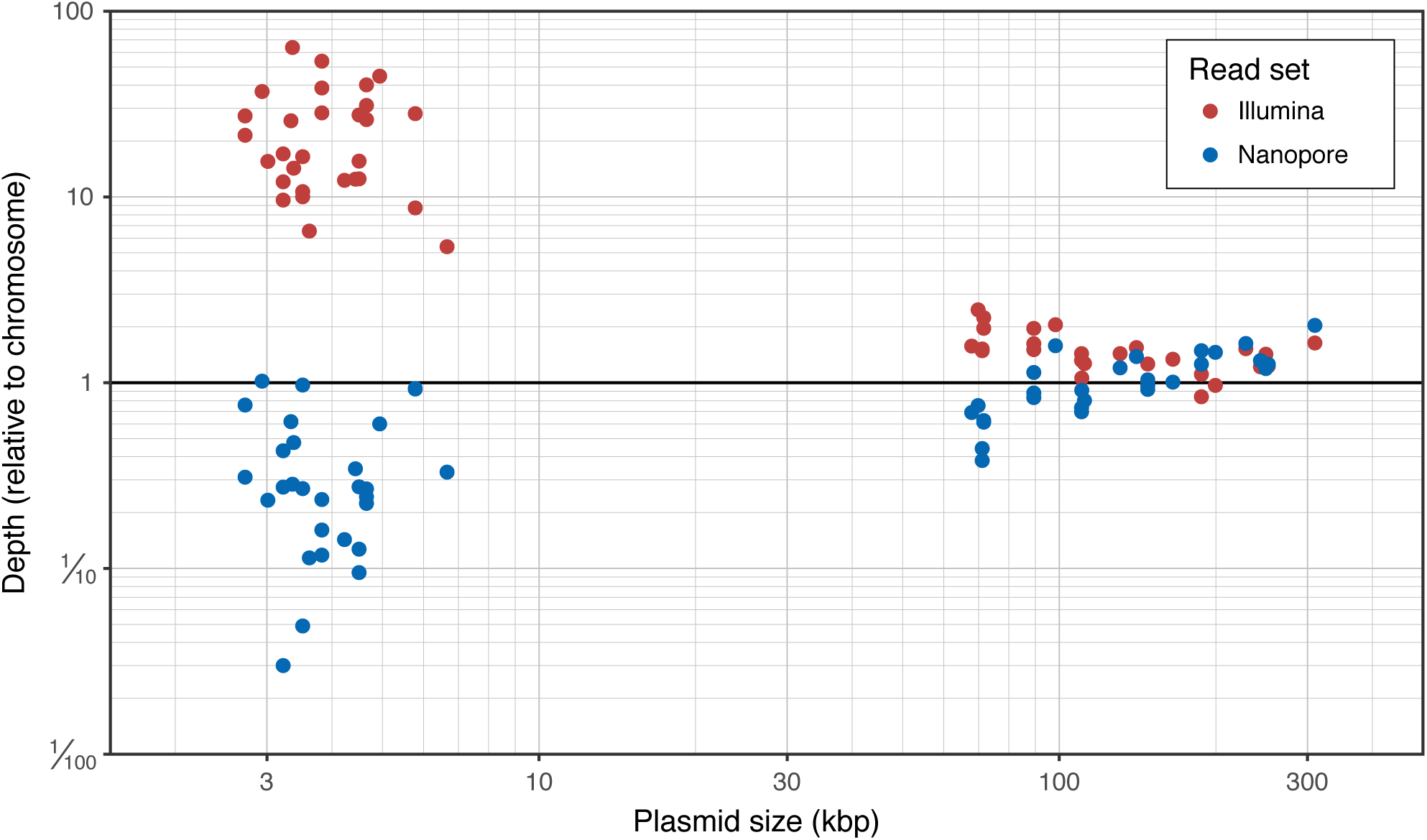
Plasmid read depths in Illumina and ONT read sets, relative to the depth of the chromosome. Note the large discrepancy in read depths of small plasmids, which are much more represented in the Illumina reads; a clear separation can be seen between small plasmids (<7 kbp) and large plasmids (>67 kbp).

**Table S1**: Characteristics of DNA libraries, sequencing results, read processing and assembly for all 12 *K. pneumoniae* samples in this study.

